# Structural basis of transcription-coupled H3K36 trimethylation by Set2 and RNAPII elongation complex in the nucleosome

**DOI:** 10.1101/2024.12.13.628464

**Authors:** Tomoya Kujirai, Haruhiko Ehara, Tomoko Ito, Masami Henmi, Shun-ichi Sekine, Hitoshi Kurumizaka

**Author notes:** Corresponding authors. (H.K.); (S.S.). These authors contributed equally to this work (Tomoya Kujirai, Haruhiko Ehara).

## Abstract

Trimethylation of the histone H3K36 residue (H3K36me3) plays an indispensable role in ensuring transcription fidelity by suppressing undesired cryptic transcription in chromatin. The H3K36me3 modification is accomplished by Set2/SETD2 during transcription elongation by the RNA polymerase II elongation complex (EC). Here we found that the Set2-mediated H3K36me3 deposition occurs primarily on the nucleosome reassembling behind the EC. Cryo-electron microscopy structures of the transcribing EC complexed with Set2 and the reassembled nucleosome revealed that Set2 is anchored by the Spt6 subunit of the EC and captures an H3 N-terminal tail of the nucleosome. Abrogation of the Set2-Spt6 interaction leads to defective transcription-coupled H3K36me3 deposition. These insights elucidate the structure-based mechanism of transcription-coupled H3K36me3 deposition in chromatin.

**One-Sentence Summary:** Cryo-EM structures of the EC-Set2-nucleosome complex reveal the mechanism of H3 Lys36 trimethylation in chromatin.

## Main Text

The nucleosome is composed of two each of four histones, H2A, H2B, H3, and H4, and 145-147 base pairs of DNA as the basic unit of chromatin in eukaryotes, and plays an essential role in the compaction of genomic DNA (*1*). The nucleosome also functions as a regulator of genome activities, such as transcription, replication, recombination, and repair (*2–4*). In transcription, the nucleosome suppresses the inappropriate production of RNA molecules by RNA polymerases, and also modulates the gene expression level by suppressing transcription processes (*3, 5*). In chromatin, transcription fidelity is ensured by the trimethylation of the Lys36 residue in the N-terminal tail of histone H3 (H3K36me3), which plays an important role in suppressing undesired cryptic transcription (*6–9*). In addition, H3K36me3 reportedly regulates DNA methylation, histone acetylation, chromatin remodeling, RNA processing, and DNA repair in the genome (*10–12*). Yeast Set2 (SETD2 in higher eukaryotes) promotes the H3K36me3 deposition during the transcription elongation process by RNA polymerase II (RNAPII) (*13–16*).

To produce nascent RNAs in chromatin, RNAPII transcribes the DNA wrapped around the histone core in nucleosomes by incrementally uncoiling the nucleosomal DNA until reaching its center (superhelical location (SHL) 0) (*17–19*). The nucleosome is then disassembled when RNAPII passes through the SHL(0) position of the nucleosome (*20*). Transcription elongation factors, Spt4/5, Elf1, Spt6, Spn1, and Paf1C, associate with RNAPII to form the RNAPII elongation complex (EC). The EC exhibits substantially enhanced activity for nucleosome transcription (*18, 20–23*). After the disassembly process, the nucleosome is readily reassembled by histone transfer to the upstream DNA behind the transcribing EC with the aid of the histone chaperone FACT (*20*). This EC/FACT-mediated nucleosome reassembly occurs on the EC upstream surface, the “cradle” formed by Spt4, Spt5, Spt6, and the Paf1C subunits, Leo1 and Rtf1, on the rim of the DNA exit of RNAPII (*20*).

The Set2 methyltransferase catalyzes the H3K36me3 deposition in the nucleosome during the transcription elongation process by the EC (*13–16*). Set2 consists of several conserved domains, including catalytic (AWS, SET, and Post-SET), AID (autoinhibitory domain), WW, CC (coiled-coil), and SRI (Set2 Rpb1-interacting) (*12, 24*) domains. The catalytic activity is reportedly regulated by the AID domain, which is modulated by the SRI domain binding to the phosphorylated C-terminal domain of RNAPII (*25–27*). Previous genetic studies suggested that the H3K36me3 deposition by Set2 is dependent on Spt6 (*28*). However, the mechanism by which Set2 specifically deposits H3K36me3 in the nucleosome in a transcription-coupled manner has remained enigmatic.

In the present study, we found that Set2 introduces H3K36me3 predominantly during the nucleosome reassembly process on the upstream side of the EC. Cryo-electron microscopy (cryo-EM) structures of the transcribing EC complexed with Set2 and the reassembled nucleosome revealed that Spt6 anchors Set2, and the Set2 catalytic domain binds a rewrapping nucleosome intermediate on the EC cradle and incorporates the N-terminal tail of the nucleosomal histone H3 within its catalytic center. These results reveal the mechanism of H3K36me3 deposition in chromatin during transcription elongation.

### Transcription elongation of the RNAPII elongation complex is required for nucleosomal H3K36me3 deposition by Set2

To study the transcription-coupled H3K36me3 deposition, we first established an *in vitro* co-transcriptional methylation assay based on the nucleosomal transcription system using the yeast *Komagataella phaffii* (formerly *K. pastoris*) EC (*20*). The EC contains RNAPII and various elongation factors, including TFIIS, Spt4/5, Elf1, Spt6, Spn1, and Paf1C (Paf1, Leo1, Ctr9, Cdc73, and Rtf1) (Fig. 1A, fig. S1), which are important for transcription elongation over the nucleosome. We prepared four nucleosome templates, termed Temp42, Temp49, Temp58, and Temp115, each containing a poly-T track at positions 42, 49, 58, and 115 base pairs from the nucleosome entry site, respectively (Fig. 1B). We then conducted the nucleosome transcription reaction by the EC with 3’-dATP instead of ATP, so that the transcription elongation would stall at the poly-T track, in the presence of Set2 and S-adenosylmethionine, a methyl donor.

**Fig. 1.**
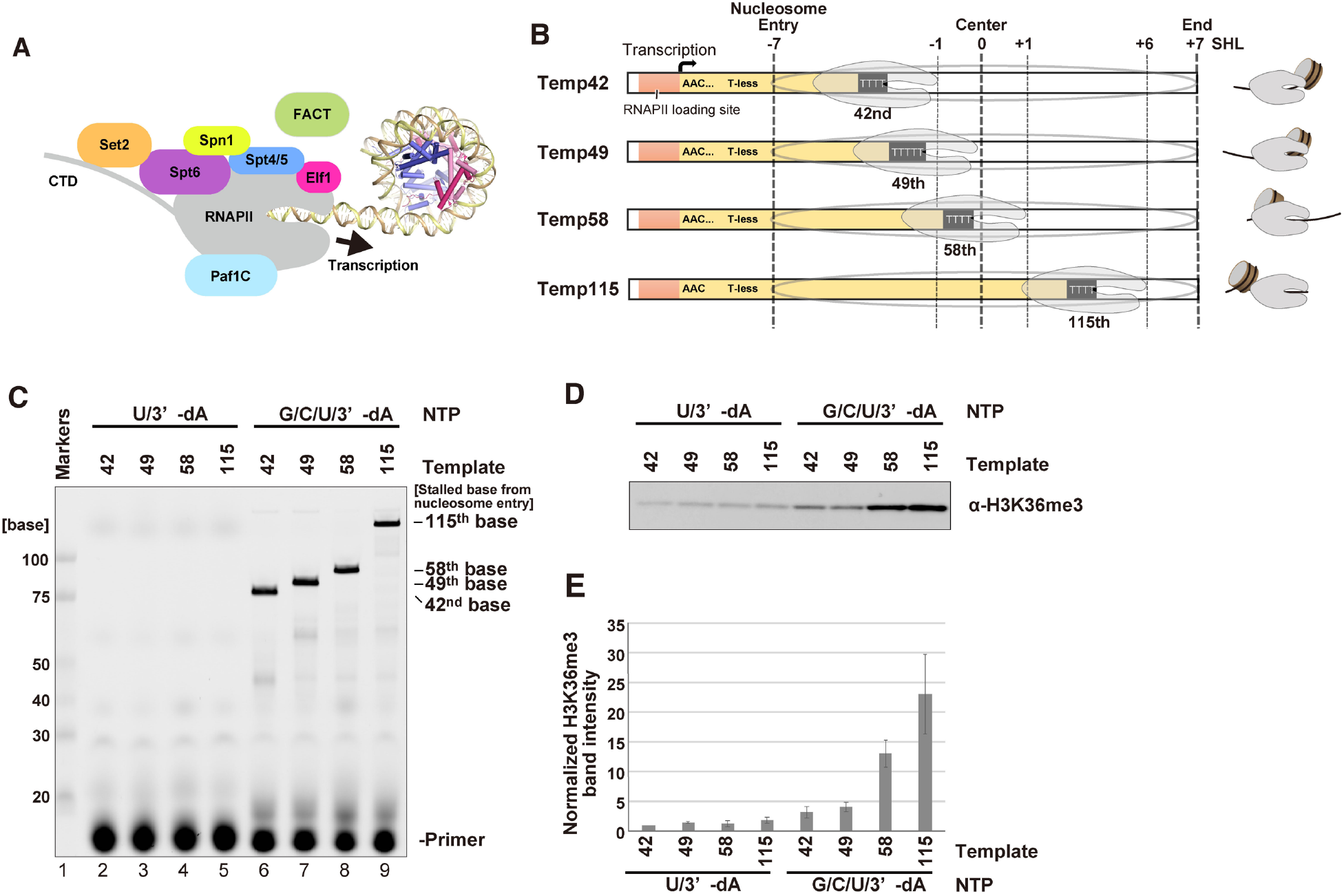
Transcription-coupled H3K36me3 deposition by Set2. (**A**) Schematic representation of *in vitro* transcription on the nucleosome template. (**B**) The nucleosome templates used in the H3K36me3 deposition assay. Cartoons depicting the relative position of a nucleosome to RNAPII are shown. (**C**) Urea PAGE analysis of the elongated RNA by EC. (**D**) Western blot of H3K36me3 deposition in different nucleosome templates. (**E**) Quantification of panel (D). The mean and SD of the relative values of the H3K36me3 signal intensities compared to that of temp42 with UTP and 3’-dATP of three independent experiments are shown.

Consequently, the transcription elongation was efficiently promoted, and the EC stalled when the RNAPII catalytic center reached the 42, 49, 58, and 115 base-pair positions, where the leading edge of RNAPII is located at the SHL(-1), SHL(0), SHL(+1), and SHL(+6) positions on the nucleosomal DNA, respectively (Fig. 1C, lanes 6-9).

In these reactions, when the transcription elongation was blocked after two bases of EC progression, by omitting GTP and CTP, a small amount of H3K36me3 was detected regardless of the template DNA utilized (Fig. 1C-E). In contrast, when the transcription was allowed to proceed into the nucleosome, a marked increase in H3K36me3 deposition was detected (Fig. 1C-E). These results indicate that the optimal H3K36 trimethylation can only occur co-transcriptionally, and the close interplay between the EC and the nucleosome is critical for the Set2 activity.

Notably, compared to the Temp42 or Temp49 substrate, the H3K36me3 deposition was substantially enhanced with the Temp58 substrate (Fig. 1D, E), and even more prominent with the Temp115 substrate. In a previous structural study, at the 42 and 49 base-pair positions, the disassembling nucleosome was observed on the downstream side of the EC (*20*). In contrast, at the 58 and 115 base-pair positions, the (sub-)nucleosome was already transferred to the upstream side of the EC. Therefore, Set2 is likely to promote the H3K36me3 deposition more efficiently on the reassembling nucleosome on the upstream side of the EC, rather than on the one remaining on the downstream side of the EC.

### Structures of the EC-nucleosome-Set2 complex stalled after passing through the nucleosomal dyad

To understand the mechanism by which Set2 preferentially promotes the H3K36me3 deposition as the nucleosome reassembles behind the EC, we performed the cryo-EM analysis of the transcribing EC stalled at the SHL(+6) position with the nucleosome template containing Temp115 DNA in the presence of Set2 (EC-nucleosome-Set2 complex). To capture the active form of the complex mediating the H3K36me3 deposition, we used the nucleosome containing H2BK120ub and H3.3K36M, in which the H2BK120 and H3.3K36 residues were replaced by a ubiquitin-conjugated cysteine residue and a methionine residue, respectively (fig. S2).

H2BK120ub reportedly stimulates H3K36me3 deposition by Set2 (*29*), and the H3.3K36M mutant, an oncogenic driver mutation, is known to irreversibly bind the catalytic center of H3K36 methyltransferases (*30–34*). The EC-nucleosome-Set2 complexes were formed by the *in vitro* nucleosome transcription reaction and fractionated by sucrose gradient ultracentrifugation in the presence of glutaraldehyde (GraFix). Cryo-EM single particle analysis yielded two structures of the EC-nucleosome-Set2 complexes, EC-nucleosome-Set2^A^ and EC-nucleosome-Set2^B^ (Fig. 2, figs. S3-8, and Tables S1-2).

**Fig. 2.**
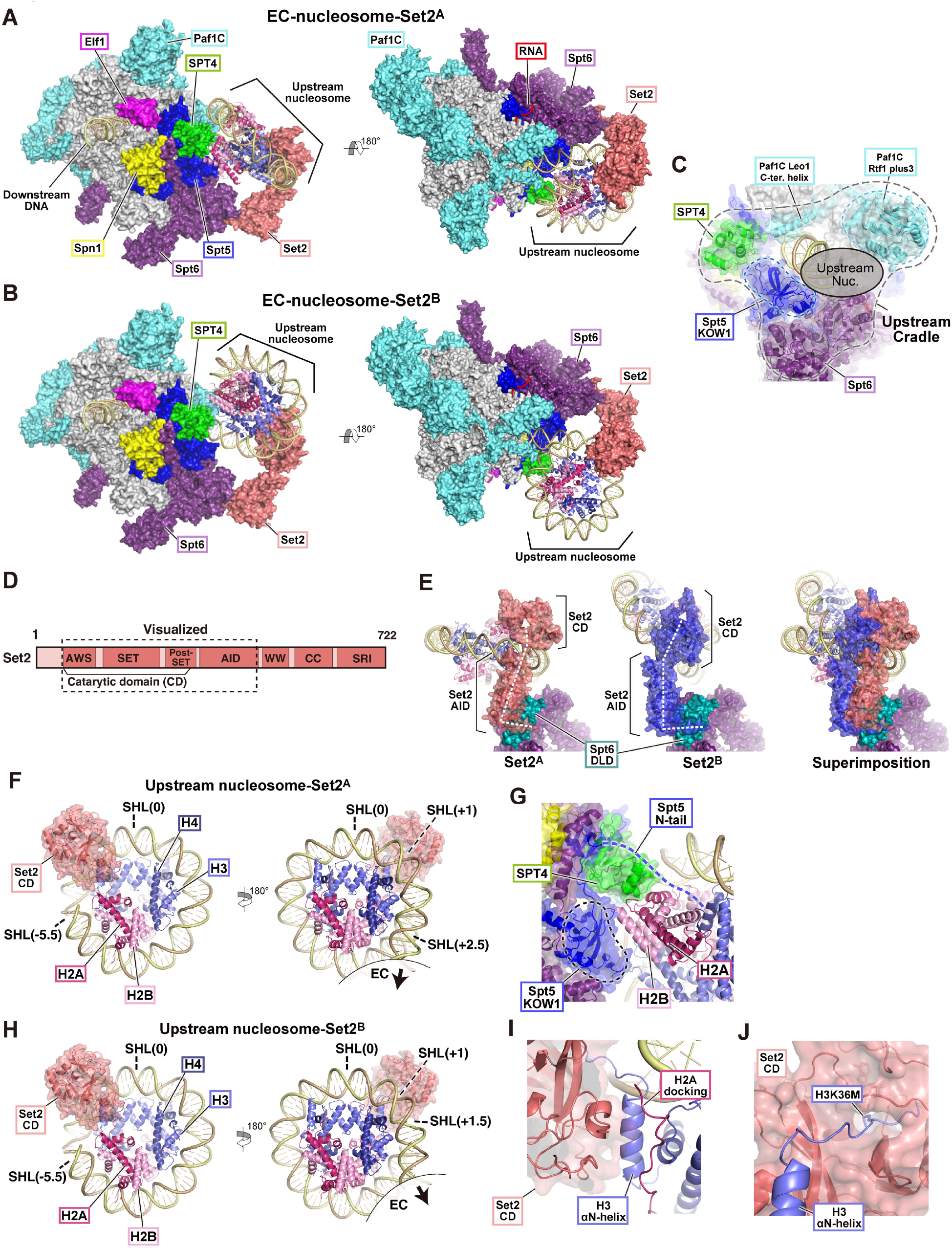
Cryo-EM structures of EC-nucleosome-Set2. (**A**) Overall structure of the EC-upstream-nucleosome-Set2^A^. (**B**) Overall structure of the EC-upstream-nucleosome-Set2^B^. The EC and Set2 structures are shown in surface models. The nucleosome structure is shown in a ribbon model. (**C**) Close-up view of the upstream surface of the EC. The elongation factors associated with RNAPII are shown in ribbon models with transparent surface models. The nucleosome and Set2 are omitted for clarity. (**D**) Domain structure of the Set2 protein. (**E**) Comparison of the Set2 structures in the EC-nucleosome-Set2^A^ and EC-nucleosome-Set2^B^ complexes. The path of Set2 from the Spt6 interaction peptide-AID-CD is shown by the dotted white line. (**F**) Structure of the nucleosome with the Set2 CD contained in the EC-nucleosome-Set2^A^. The CD is shown in a ribbon model with a transparent surface model. (**G**) Close-up view of the upstream surface of the EC – nucleosome contacts. (**H**) The nucleosome structure contained in the EC-nucleosome-Set2^B^. (**I**) Close-up view of the Set2 CD, H3 αN-helix, and H2A docking domain contacts. (**J**) Close-up view of the H3 N-terminal tail and the Set2 CD. The H3 Met36 residue is shown in a stick model.

In the EC-nucleosome-Set2 structures, we successfully observed Set2 associated with the reassembled nucleosome behind the EC, in which the elongation factors Spt4, Spt5, Spt6, Spn1, Elf1, and Paf1C are visualized together with RNAPII (Fig. 2A, B). Spt4, the Spt5 KOW1 domain, Spt6, the Rtf1 Plus3 domain and the Leo1 C-terminal helix of Paf1C form the “cradle” on the upstream face of the EC, which supports nucleosome reassembly after histone transfer (Fig. 2C) (*20*). Overall, the assembly of the transcription elongation factors on RNAPII is essentially identical between the EC-nucleosome-Set2^A^ and EC-nucleosome-Set2^B^ structures and is similar to that of the previously reported EC structure (*20*). In these structures, the Set2 catalytic domain (CD) and AID domain are visible (Fig. 2D). Set2 is anchored by Spt6 and binds to the reassembling nucleosome intermediate on the cradle (Fig. 2A, B). These two EC-nucleosome-Set2 complexes accommodate nucleosomes in different orientations with the same Set2 CD-nucleosome interactions, reflecting the flexibility of the Set2 structure. In fact, in the EC-nucleosome-Set2^A^ and EC-nucleosome-Set2^B^ structures, the Set2 structures differ in the relative orientation between CD and AID, because of the flexibility between the two domains (Fig. 2E). In addition, the Set AID orientation relative to Spt6 differs between the EC-nucleosome-Set2^A^ and EC-nucleosome-Set2^B^ structures, indicating that the Spt6-binding interface of Set2 is also flexible (Fig. 2E).

In the EC-nucleosome-Set2^A^ structure, ∼90 base-pairs of DNA [SHL(-5)-SHL(+2.5)] are wrapped around the histone octamer (Fig. 2F). Consistent with our previous report, in the reassembled nucleosome, Spt4 and the Spt5 KOW1 domain directly contact the exposed H2A-H2B and H3-H4 surfaces of the nucleosome on the cradle (Fig. 2G). In addition, a weak cryo-EM density, probably corresponding to the Spt5 N-terminal tail, is observed on the exposed H2A-H2B surface. This finding supports the previous report that the Spt5 N-terminal tail binds the histone complex and promotes nucleosome reassembly (*35*) (Fig. 2G). In the EC-nucleosome-Set2^B^ structure, ∼80 base-pairs of DNA [SHL(-5)-SHL(+1.5)] are wrapped around the histone octamer (Fig. 2H). The orientation of the nucleosome relative to the RNAPII is different compared to that of the EC-nucleosome-Set2^A^ structure, although the exposed H2A-H2B surfaces contact the EC cradle in both complexes (Fig. 2A, B). In both nucleosome structures, the Set2 CD binds the nucleosomal DNA around the SHL(+1) position and interacts with the H3 αN-helix and H2A C-terminal docking domain, exposed to the solvent due to incomplete DNA rewrapping around the SHL(-7)-SHL(-6) region (Fig. 2F, 2H, 2I). The K36M residue in the H3 N-terminal tail is incorporated into the catalytic center of the Set2 CD (Fig. 2J). This Set2 CD binding to the nucleosome is consistent with the previous studies without RNAPII transcription (*29, 31*). The ubiquitin moiety of H2BK120C was not visible in the present structures.

### Set2-Spt6 interaction is critical for transcription-coupled H3K36me3 deposition

Interestingly, we found that the region of amino acid residues 453-457 (YKIPK) of the Set2 AID directly binds the Spt6 DLD (death-like domain) (Fig. 3A, B). Particularly, the Set2 K454 and K457 residues may interact with the acidic region of the Spt6 DLD via electrostatic interactions. In Set2, I455 hydrophobically interacts with the Spt6 M1058, L1103, and Y1108 residues (Fig. 3B). The *K. phaffii* Set2 YKIPK residues in the Spt6-binding motif are mostly conserved in other eukaryotes, including yeast, worm, fly, mouse, and human (Fig. 3C). This Set2 AID domain-Spt6 interaction tethers the Set2 CD near the cradle of the EC and promotes the CD binding to the nucleosomal DNA at the SHL(+1) position and the H3 N-terminal tail (Fig. 3A).

**Fig. 3.**
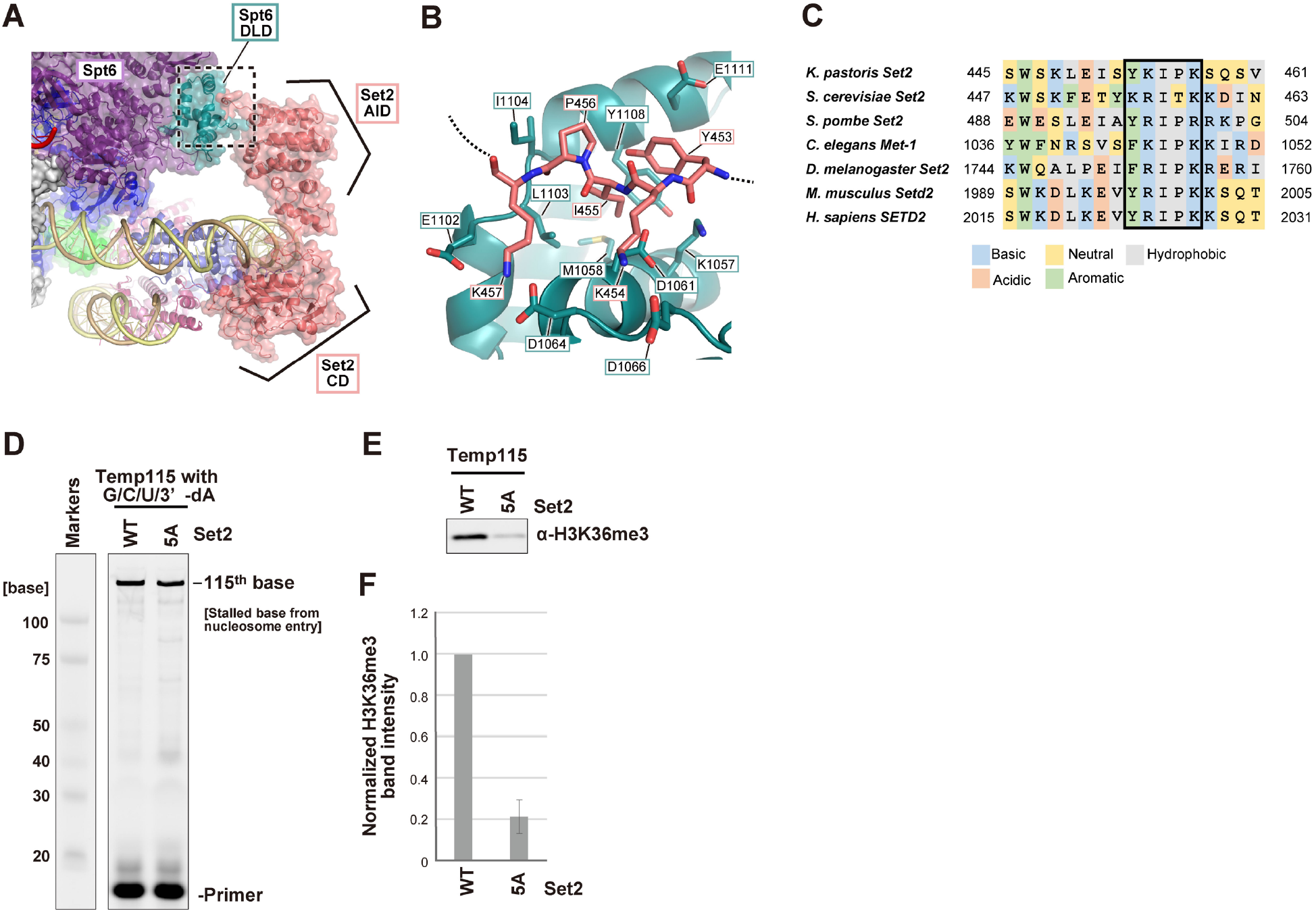
Interface between Set2 and Spt6. (**A**) Close-up view of the Spt6-Set2-upstream nucleosome in the EC-nucleosome-Set2^A^. (**B**) Close-up view of the Spt6-Set2 interaction. This view corresponds to the dotted rectangle in panel (A). Nitrogen and oxygen are colored blue and red, respectively. The Set2 453-457 residues and Spt6 residues, which may interact with Set2, are shown in stick models. (**C**) Sequence alignment of Set2 proteins from various model organisms. The YKIPK motif is enclosed by a black rectangle. (**D**) Urea PAGE analysis of the elongated RNA in the transcription assay with the Set2 wild-type (WT) or 5A mutant. The transcription reaction was conducted using the temp115 nucleosome with GTP, CTP, UTP, and 3’-dATP. (**E**) Western blot of H3K36me3. The samples shown in panel (D) were analyzed. (**F**) Quantification of the H3K36me3 band intensity shown in panel (E). The mean and SD of the relative values of the H3K36me3 signal intensities compared to that of the Set2 wild-type (WT) in three independent experiments are shown.

To study the functional relevance of the Set2-Spt6 interaction, we prepared the Set2 mutant, Set2^5A^, in which five residues of the Spt6 binding region of Set2, 453-457 (YKIPK), are replaced by alanine. We then performed the EC transcription-coupled H3K36me3 deposition assay with the Temp115 nucleosome. The Set2^5A^ mutation did not affect the transcription efficiency of the EC with the Temp115 nucleosome (Fig. 3D). In the absence of the EC, the Set2^5A^ mutant mediated H3K36me3 deposition as effectively as wild-type Set2 (fig. S9).

However, the Set2^5A^ mutant was substantially defective in the transcription-coupled H3K36me3 deposition (Fig. 3E, F). These results indicate that the Set2 AID domain-Spt6 interaction plays a critical role in the transcription-coupled H3K36me3 deposition.

### Set2 binding is incompatible with FACT binding and a complete nucleosome

Previously, we reported that the EC stalled at the position 58 bp from the nucleosome entry (EC58^oct^) contains a subnucleosome structure on the cradle, in which ∼75 base-pairs of DNA [SHL(-7)-SHL(+0.5)] are wrapped around the histone octamer complexed with FACT (*20*). In EC58^oct^, FACT contacts the nucleosomal DNA SHL(-1)-SHL(0) region, and the HA1 helix of the FACT Spt16 subunit binds the exposed surface of H3-H4 corresponding to the SHL(+1) region, instead of DNA (Fig. 4A). We found that Set2 and FACT binding are mutually exclusive, because Set2 would cause a steric clash with the DNA around SHL(+1) in the subnucleosome intermediate complexed with FACT (Fig. 4E). Given that Set2 binds the nucleosome in which the nucleosomal DNA SHL(+1) is rewrapped (Fig. 4B, C), Set2 may be able to bind the nucleosome after FACT dissociates from the H3-H4 surface corresponding to the SHL(+1) DNA region. Consistent with this idea, the H3K36me3 deposition is significantly enhanced in the Temp115 nucleosome, which may have released FACT from the nucleosome, as compared to the Temp58 nucleosome (Fig. 1D, E).

**Fig. 4.**
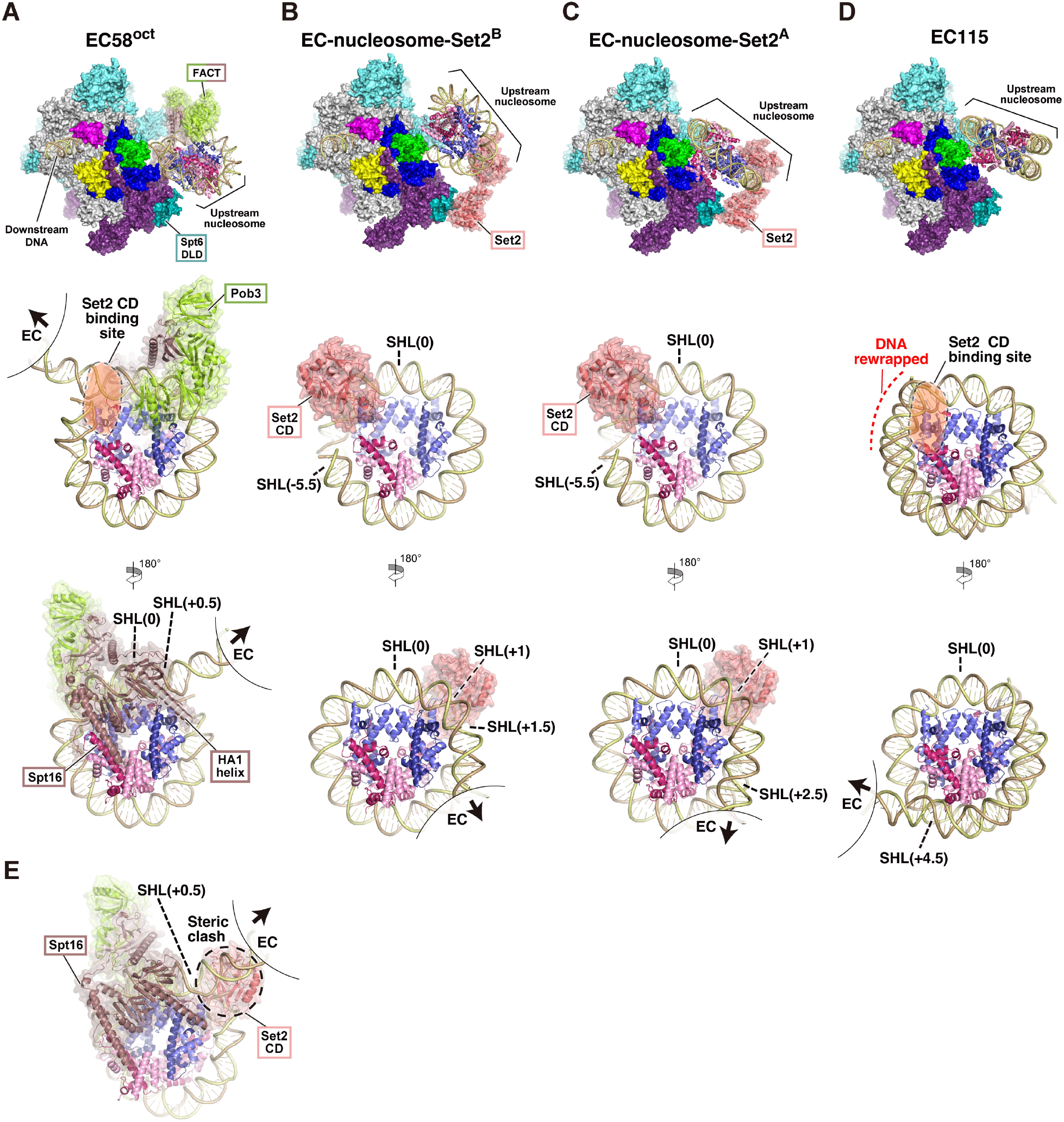
Structural comparison of the reassembling nucleosomes. (**A-D**) The overall and nucleosome structures contained in EC58oct (**A**), EC-nucleosome-Set2^B^ (**B**), EC-nucleosome-Set2^A^ (**C**), and EC115 (**D**). The CD binding site is shown. The FACT and Set2 CD structures are shown in ribbon models with a transparent surface model. The HA1 helix of the FACT Spt16 is highlighted. (**E**) Superimposed view of the nucleosome-FACT of EC58^oct^ with the Set2 CD of EC-nucleosome-Set2^A^. The Set2 CD clashes with the unwrapped nucleosomal DNA SHL(+1) of EC58^oct^.

We also compared the current EC-nucleosome-Set2 structures with the previously reported EC115 structure (*20*). In EC115, ∼120 base pairs of DNA [SHL(-7)-SHL(+4)] are rewrapped around the histone octamer (Fig. 4D). The rewrapped nucleosomal DNA end around the SHL(-7)-SHL(-6) region conceals the H3 αN-helix. In contrast, in the EC-nucleosome-Set2 complex, the SHL(-7)-SHL(-6) region is stripped from the histone surfaces and the Set2 CD contacts the exposed H3 αN-helix (Fig. 4B, C). Therefore, the nucleosome in EC115 is incompatible with Set2 CD binding. This suggests that Set2 may need to complete H3K36me3 deposition before rewrapping the SHL(-7)-SHL(-6) region, although we cannot exclude the possibility that Set2 unwraps the SHL(-7)-SHL(-6) region by itself, albeit less efficiently.

## Discussion

In the present study, we demonstrated that the nucleosomal H3K36me3 deposition by Set2 requires transcription elongation through the nucleosome by the EC. The cryo-EM structures of the EC-nucleosome-Set2 complexes capture the H3K36me3 deposition during nucleosome reassembly and reveal the crucial interactions between Set2 and Spt6 and between Set2 and the reassembling nucleosome intermediate.

We discovered that the H3K36me3 deposition by Set2 is significantly enhanced when the EC passes through the SHL(0) position of the nucleosome. This finding suggests that Set2 primarily promotes H3K36me3 deposition during the nucleosome reassembly process on the upstream side of the EC. In the EC-nucleosome-Set2 structures, we found that the Set2 AID binds to the Spt6 subunit of the transcribing EC via the Spt6-binding motif, which is conserved from yeast to human (Fig. 3C). Our mutational analysis of the Spt6-binding motif of Set2 revealed that the Set2-Spt6 binding plays an important role in the EC transcription-coupled H3K36me3 deposition (Fig. 3D, E). The binding between Set2 and Spt6 optimally positions the Set2 CD for interactions with the nucleosomal DNA and the H3 αN-helix region, allowing efficient H3K36me3 deposition during nucleosome reassembly by the EC.

The structures of nucleosomes complexed with the CD of yeast Set2 and its human homologue SETD2 have been reported (*29, 31*). These previous studies revealed that the CD of Set2 and SETD2 binds the nucleosomal DNA and the αN-helix of histone H3. The interaction between the CD-H3 αN-helix plays an essential role in incorporating the target Lys36 residue of H3 into the catalytic center of the CD. To accomplish this, the CD binds the nucleosomal DNA at the SHL(+1) position, and unwraps the DNA around the nucleosomal DNA end, exposing the H3 αN-helix (*29, 31*). Consistently, in the present study, we found that the CD of Set2 contacts the rewrapped nucleosomal DNA at the SHL(+1) position in the EC-nucleosome Set2 complexes. However, during the nucleosome reassembly process, the H3 αN-helix should be exposed to the solvent to allow efficient CD binding (Fig. 2G, I). This explains why transcription is required for the efficient deposition of H3K36me3.

Based on these results and the previous structures (EC58^oct^ and EC115), we propose a model of transcription-coupled H3K36me3 deposition (Fig. 5). When the EC passes through the dyad (SHL(0)) of a nucleosome, histones are transferred from the downstream side of the EC to the upstream side, where nucleosome reassembly begins. In the early stage of the reassembly (EC58^oct^), the histones in the subnucleosome intermediate are bound by FACT, and the DNA is still unwrapped to SHL(+1), which are both incompatible with Set2 binding. Although the Set2 AID is bound to Spt6, the Set2 CD is unable to bind to the nucleosome and is in a “standby” mode. As the EC advances, the DNA would gradually rewrap the histone surfaces, facilitating FACT detachment from the histones. When the DNA is rewrapped around SHL(+1), the Set2 CD binds to the nucleosome to catalyze H3K36me3 deposition. After the reaction, the Set2 CD probably dissociates from the nucleosome, and further DNA rewrapping would complete the nucleosome formation with the H3K36me3 modification.

**Fig. 5.**
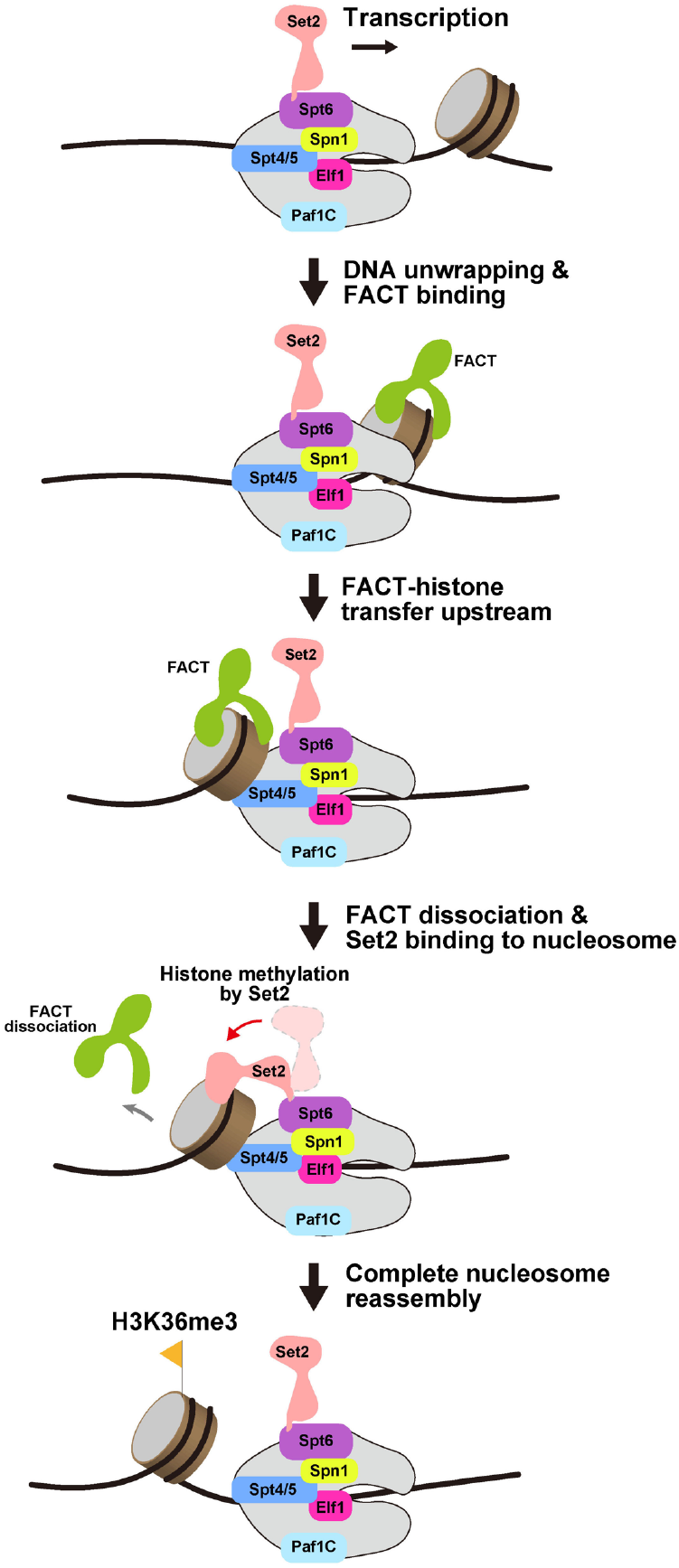
Model of H3K36me3 deposition during transcription through the nucleosome. See discussion.

Set2/SETD2 (mammalian homologue) reportedly mediates the H3K36me3 deposition in the nucleosome without RNAPII and elongation factors *in vitro* (*6, 29, 31*). Consistently, our nucleosomal H3K36me3 deposition assay also detected the Set2 methyltransferase activity in the absence of the EC (fig. S7D). However, the H3K36me3 deposition was barely detectable, when H3K36 methylation was examined in the presence of RNAPII and elongation factors, but limited nucleotide triphosphates (U/3’-dATP, Fig. 1D,E). This discrepancy may be explained by the possibility that Set2 is sequestered from the downstream nucleosome by the EC, which is distant from the nucleosome. Efficient H3K36me3 deposition may require transcription elongation that accompanies nucleosome reassembly in proximity to Set2 tethered by Spt6.

A previous genetic study reported that yeast cells harboring the spt6 mutant, *spt6-1004*, which lacks the HhH domain, are substantially defective in H3K36me3 deposition (*28*). The Spt6 HhH region interacts with not only the Spt5 KOW1 domain but also the Spt6 DLD domain, which is the Set2 AID domain binding site (fig. S10). Therefore, the deletion of the Spt6 HhH domain may allosterically affect the structure or location of the Spt6 DLD domain in the EC structure, leading to inappropriate Set2 positioning.

SETD2 is implicated in tumorigenesis. SETD2 mutations and loss of function have been found in tumors (*36, 37*). Interestingly, five mutations in the region of human SETD2 amino acid residues 2023-2027 (YRIPK), the conserved SPT6 interaction site, have been found in patients with carcinoma (fig. S11). Among these mutations, four were the SETD2 R2024Q mutation. The *K. phaffii* Set2 K454Q mutation, which corresponds to the human SETD2 R2024Q mutation, may affect the interaction between Set2 and Spt6, and consequently reduce H3K36me3 deposition. These findings highlight the importance of the interaction between Spt6/SPT6 and Set2/SETD2 for proper H3K36me3 deposition and gene regulation.

## Acknowledgements

We thank Y. Iikura, Y. Takeda, J. Kato (The University of Tokyo), M. Aoki, and M. Goto (RIKEN) for their assistance. The cryo-EM experiments were performed with the Krios G4 microscope at the RIKEN Yokohama cryo-EM facility.

## Funding

Japan Society for the Promotion of Science KAKENHI JP20H05690 (SS, TK)

Japan Society for the Promotion of Science KAKENHI JP24H00062 (SS, TK)

Japan Society for the Promotion of Science KAKENHI JP20H03201 (HE, TK)

Japan Society for the Promotion of Science KAKENHI JP23K17392 (TK)

Japan Society for the Promotion of Science KAKENHI JP22K15033 (TK)

Japan Society for the Promotion of Science KAKENHI JP24H02328 (HK)

Japan Society for the Promotion of Science KAKENHI JP23H05475 (HK)

Japan Society for the Promotion of Science KAKENHI JP20H05906 (SS)

Japan Science and Technology Agency ERATO JPMJER1901 (HK)

Research Support Project for Life Science and Drug Discovery (BINDS) from AMED JP23ama121009 (HK)

Daiichi Sankyo Foundation for Life Science (SS)

## Author contributions

Conceptualization: TK, HE, HK, SS

Methodology: TK, HE, TI, MH

Investigation: TK, HE, TI

Visualization: TK, HE, HK, SS

Funding acquisition: TK, HE, SS, HK

Project administration: TK, HE, SS, HK

Supervision: HK, SS

Writing – original draft: TK, HE, SS, HK

Writing – review & editing: TK, HE, SS, HK

## Data and materials availability

The cryo-EM maps and coordinates have been deposited in the Electron Microscopy Data Bank (EMDB) and Protein Data Bank (PDB), respectively, with accession codes: EMD-XXXXX and XXXX (EC-nucleosome-Set2^A^), and EMD-xxxx and xxxx (EC-nucleosome-Set2^B^).

