## Supplementary_materials for "Structural basis of transcription-coupled H3K36 trimethylation by Set2 and RNAPII elongation complex in the nucleosome"

**The PDF file includes:**

Materials and Methods
Figs. S1 to S11
Table S1-2
References 38-57

### Materials and Methods

Protein preparation

*K. phaffii* RNAP II, Spt4/5, Elf1, Spn1, Spt6, Paf1C, FACT, human P-TEFb, human histones H2A, H2B (wild-type and K120C mutant), H3.1, H3.3 (wild-type and K36M mutant), and H4 were purified as described previously (*20*, *38–40*). The histone octamers, except for that containing H2BK120Cub, were prepared as described previously (*40*).

H2BK120Cub, in which ubiquitin is conjugated to Cys120 of the H2B protein, was prepared as described previously with modifications (*41*). The hexahistidine (His6)-tagged ubiquitin was purified as described previously. The H2BK120C and His6-ubiquitin powders were separately dissolved to 10 mg/ml in denaturing buffer adjusted to pH 8.5 (50 mM sodium tetraborate and 6 M urea). The resulting protein solutions were mixed in a 1:1 molar ratio, and then 5 mM TCEP was added. After rotation at room temperature for 30 minutes, the sample was cooled on ice. Then 0.44 mM DCA (1.3-dichloroacetone) dissolved in DMF (N,N-dimethylformamide) was added to crosslink the histone and ubiquitin, and the solution was incubated on ice for 30 minutes. Afterward, 5 mM 2-mercaptoethanol was added to quench the crosslinking reaction. The cross-linked sample was dialyzed against Ni wash buffer [50 mM Tris-HCl (pH 8.0), 500 mM NaCl, 6 M urea, 5% glycerol, and 5 mM imidazole]. The dialyzed sample was purified using Ni-NTA column chromatography, and eluted by Ni wash buffer containing 500 mM imidazole. The resulting sample contains ubiquitin monomer, ubiquitin dimer, and H2B-ubiquitin. This sample was dialyzed against MonoS wash buffer [20 mM KOAc (pH 5.2), 200 mM NaCl, 6M urea, 1 mM EDTA, and 5 mM 2-mercaptoethanol], and purified by MonoS cation exchange column chromatography, with elution by MonoS wash buffer containing 900 mM NaCl. The fractions containing H2B-ubiquitin were collected. The purified H2B-ubiquitin was dialyzed against distilled water and stored at -80°C. For the preparation of the histone octamer containing H2BK120ub, H2A, H2BK120Cub, H3.3, and H4 were dissolved in denaturing buffer [20 mM Tris-HCl (pH 7.5), 7 M guanidine-HCl, and 20 mM 2-mercaptoethanol] to a total protein concentration of 1.0 mg/ml. The sample was refolded during dialysis against refolding buffer [10 mM Tris-HCl (pH 7.5), 2 M NaCl, 1 mM EDTA, and 5 mM 2-mercaptoethanol], purified by Superdex200 gel filtration chromatography, flash-frozen in liquid nitrogen, and stored at -80°C.

Mouse casein kinase 2 (CK2) was purified as described previously with modifications (*42*). The DNA fragments encoding the CK2α and β subunits were inserted into the pRSFDuet vector. The CK2 α and β subunits were co-expressed as recombinant proteins in *E. coli* BL21 (DE3) RIL cells. The His6 tag was fused to the CK2α subunit. The cultured cells were harvested by centrifugation, suspended in lysis buffer [50 mM Tris-HCl (pH 7.5), 500 mM NaCl, 1 mM PMSF, and 20 mM imidazole], and disrupted by sonication. The disrupted cells were centrifuged, and the His6-tagged CK2α - β proteins contained in the supernatant were purified by Ni-NTA column chromatography. The Ni beads were washed with lysis buffer, followed by an additional wash with Ni wash buffer [50 mM Tris-HCl (pH 7.5), 100 mM NaCl, 1 mM PMSF, and 20 mM imidazole]. The CK2α - β protein was eluted with Ni wash buffer containing 300 mM imidazole. The resulting protein was purified by HiTrap Heparin chromatography (Cytiva) using heparin wash buffer [20 mM Tris-HCl (pH 7.5), 100 mM NaCl, 1 mM EDTA, and 2 mM 2-mercaptoethanol] and eluted with heparin wash buffer containing 1 M NaCl. The resulting protein was further purified on a Superose 6 Increase gel filtration column using sec buffer [20 mM Tris-HCl (pH 7.5), 1 M NaCl]. The fractions containing the CK2α protein or CK2α - β complex were separately collected. The CK2α protein was dialyzed against dialysis buffer [20 mM HEPES-NaOH (pH 7.5), 300 mM NaCl, 0.5 mM EDTA, 5% glycerol, 1 mM DTT], flash-frozen in liquid nitrogen, and stored at -80°C.

*K. phaffii* Set2 for the cryo-EM analysis was expressed as an HRV-3C cleavable, N-terminally His6-tagged protein in *Escherichia coli*. The cells were first resuspended in buffer A [20 mM Tris-HCl (pH 8.0), 700 mM NaCl, and 10 mM 2-mercaptoethanol] supplemented with 0.1 mM phenylmethylsulfonyl fluoride (PMSF), and then disrupted by sonication. The lysate was cleared by centrifugation, and applied to a Ni Sepharose 6 Fast Flow column (Cytiva). The column was washed serially with buffer A supplemented with 20 mM imidazole and buffer B150 [20 mM Tris-HCl (pH 8.0), 150 mM NaCl, 10 mM 2-mercaptoethanol, and 20 mM imidazole]. The column was treated overnight with HRV-3C protease to cleave the tag. The protein was eluted with buffer B150, and then diluted two-fold using buffer C [20 mM Hepes-KOH (pH 7.5), 0.1 μM zinc acetate, 0.1 mM tris(2-carboxyethyl)phosphine hydrochloride (TCEP-HCl), and 5% glycerol]. The protein was further purified by Resource Q anion-exchange column chromatography (Cytiva), using a linear gradient of buffer C to buffer D [20 mM HEPES-KOH (pH 7.5) and 2 M potassium acetate]. Fractions containing Set2 were collected, and then concentrated and buffer-exchanged to buffer E [20 mM HEPES-KOH (pH 7.5), 150 mM potassium acetate, 0.1 μM zinc acetate, 0.1 mM TCEP-HCl, and 5% glycerol], using an Amicon Ultra filter (Millipore). *K. phaffii* Set2 and its mutant for the assays were also expressed as HRV-3C cleavable, N-terminally His6-tagged proteins in *Escherichia coli*, and purified using an improved protocol with better yields. The cells were resuspended in buffer A and disrupted by sonication. The lysate was cleared by centrifugation, and applied to a Ni Sepharose 6 Fast Flow column (Cytiva). The column was washed serially with buffer A supplemented with 20 mM imidazole and buffer B300 [20 mM Tris-HCl (pH 8.0), 300 mM NaCl, 10 mM 2-mercaptoethanol, and 20 mM imidazole], and then treated overnight with HRV-3C protease. The protein was eluted with buffer B300, diluted two-fold using buffer C, and then further purified by Resource S cation-exchange column chromatography (Cytiva), using a linear gradient of buffer C to buffer D. Fractions containing Set2 were concentrated and buffer-exchanged to buffer E, using an Amicon Ultra filter (Millipore).

Preparation of template DNAs

All nucleosomal template DNA sequences were designed based on the Widom601 DNA sequence (*43*). The temp49 and temp115 DNA fragments were designed and purified as described previously (*20*). The DNA sequences of temp42 and temp58 were designed as described below. Briefly, the DNA fragments were amplified by PCR, cleaved by *Bgl*I, and purified by non-denaturing polyacrylamide electrophoresis (PAGE) using a Prep Cell (Bio-Rad) or Superose 6 Increase gel-filtration column chromatography (Cytiva). The purified DNA fragments were concentrated by an Amicon Ultra 3K centrifugal concentrator (Millipore) and stored at -20°C. The DNA sequences of temp42 and temp58 are as follows:

Temp42

non-template strand: 5´-
TGGCCGTTTTCGTTGTTTTTTTCTGTCTCGTGCCTGGTGTCTTGGGTGTAAAACCCTTG
GCGGTTAAAACGCGGGGGACAGCGCGTACGTGCGTTTAAGCGGTGCTAGAGCTGTC
TACGACCAATTGAGCGGCCTCGGCACCGGGATTCTGAT-3´;

template strand: 5′-
ATCAGAATCCCGGTGCCGAGGCCGCTCAATTGGTCGTAGACAGCTCTAGCACCGCTT
AAACGCACGTACGCGCTGTCCCCCGCGTTTAACCGCCAAGGGTTTACACCCAAGA
CACCAGGCACGAGACAGAAAAAAACAACGAAAACGGCCACCA-3′.

Temp58
non-template strand: 5′-
TGGCCGTTTTCGTTGTTTTTTTCTGTCTCGTGCCTGGTGTCTTGGGTGTTTTCCCCTTG
GCGGTTAAACGCGGGGGACAGCGCGTACGTGCGTTTAAGCGGTGCTAGAGCTGTC
TACGACCAATTGAGCGGCCTCGGCACCGGGATTCTGAT-3′;

template strand: 5′-
ATCAGAATCCCGGTGCCGAGGCCGCTCAATTGGTCGTAGACAGCTCTAGCACCGCTT
AAACGCACGTACGCGCTGTCCCCCGCGTTTAACCGCCAAGGGGAAAACACCCAAG
ACACCAGGCACGAGACAGAAAAAAACAACGAAAACGGCCACCA-3′;

Nucleosome preparation

The template nucleosomes containing H2A, H2B (wild-type or K120Cub), H3.3 (wild-type or K36M mutant), and H4 were reconstituted by the salt dialysis method and purified as described previously (*40*). After dialysis, the short double-stranded DNA fragment (*17*) was ligated to the sticky end of the nucleosomal DNA. For the nucleosome containing H2BK120Cub, the His6 tag fused to the ubiquitin was cleaved by TEV protease. The resulting nucleosomes were then purified by non-denaturing polyacrylamide gel electrophoresis using a Prep Cell (Bio-Rad). The nucleosome containing H2A, H2B, H3.1, H4, and the Widom601 193 base-pair DNA (*44*) was prepared as described previously (*20*, *40*). The purified nucleosomes were flash-frozen in liquid nitrogen and stored at -80°C.

Transcription-coupled H3K36me3 deposition assay

The transcription reaction was performed by mixing the indicated template nucleosomes, RNAPII, TFIIS, Spt4/5, Elf1, Paf1C, Spt6, Spn1, FACT, P-TEFb, CK2$\alpha$, Set2, and a DY647 fluorescently labeled RNA primer (Dharmacon) in 27 μL of reaction solution, containing a UTP and 3′-dATP combination or a UTP, GTP, CTP, and 3′-dATP combination, and incubated at 37°C for 30 minutes. In this step, the RNAPII elongation complex proceeded and stalled at the first C position with the UTP and 3′-dATP combination or the first T position with the UTP, GTP, CTP, and 3′-dATP combination. CK2 phosphorylates Spt5, Spt6, and FACT, promoting Spt6 binding to Spn1 and FACT binding to histones (*45–48*). P-TEFb phosphorylates the RNAPII C-terminal domain, Spt5, and Paf1C, promoting the EC assembly (*49*, *50*). After this incubation, 3 μL of S-adenosylmethionine (SAM) was added to the reaction solution, which was incubated for 1 hour at 37°C for histone methylation. The resulting reaction mixture contains 0.1 μM template nucleosome, 0.1 μM RNAPII, 0.1 μM TFIIS, 0.4 μM Spt4/5, 1.0 μM Elf1, 0.2 μM RNA, 0.27 μM CK2$\alpha$, 0.1 μM P-TEFb, 0.4 μM Spt6, 0.5 μM Spn1, 0.4 μM Paf1C, 0.5 μM FACT, and 1.0 μM Set2, in 31 mM HEPES-KOH (pH 7.5) buffer containing 68 mM KOAc, 0.4 μM $Zn(OAc)_2$, 0.04 mM TCEP, 2.5 % glycerol, 15 mM NaCl, 0.2 mM DTT, 5 mM $MgCl_2$, 0.02 mM EDTA, 0.1 mM SAM, and NTPs (0.4 mM UTP and 0.4 mM 3′-dATP, or 0.4 mM UTP, 0.4 mM GTP, 0.4 mM CTP, and 0.4 mM 3′-dATP).

For the urea-PAGE analysis of RNA, 2 μL of the reaction mixture was mixed with 1 μL of stop solution (100 mM Tris-HCl (pH 7.5), 1.0 mg/ml Proteinase K (Roche), 150 mM EDTA, and 4 M urea) and incubated at room temperature for 10 min, followed by adding 12 μL of HiDi formamide (ThermoFisher) and heating at 95°C for 5 minutes. The resulting RNA was analyzed by 10% denaturing urea PAGE. Blue-colored RNA markers (Biodynamics Laboratory) were used. The DY647 fluorescent signal was detected by an Amersham Typhoon imager (Cytiva) through a glass plate. The images were adjusted by ImageJ (*51*) with linear contrast enhancement.

For the western blot analysis of H3K36me3, 24 μL of the reaction mixture was mixed with 8 μL of 4x SDS buffer, and then heated at 95°C for 1 minute. A 15 μL portion of the resulting sample was fractionated by SDS-PAGE (Nacalai Tesque, Extra PAGE One Precast Gel 10-20%, cat. No. 13068-24). PageRuler Plus (ThermoFisher, cat. No. 26619) protein markers were used. After electrophoresis, the proteins in the gel were transferred to a membrane (Cytiva, Amersham Protram 0.1 μm NC cat. No. 10600005 or Amersham Protran Premium NC 0.2 μm, cat. No. 10600009). The membrane was blocked with Blocking one-P (Nacalai Tesque, cat. No. 05999-84) for 20 minutes. After washing with TBS-T three times, the membrane was incubated with a mouse monoclonal anti-H3K36me3 antibody (Active Motif, cat. No. 61022, 1:1,000) overnight at 4°C. The antibody was diluted with Can Get Signal Solution 1 (TOYOBO, cat. No. NKB-201). After the membrane was washed with TBS-T three times, it was incubated with a goat anti-mouse IgG conjugated with Cy3 (Jackson Immuno Research Laboratories, Inc., cat. No. 115-165-003, 1:100) for 1 hour. This antibody was diluted with Can Get Signal Solution 2 (TOYOBO, cat. No. NKB-301). After washing the membrane four times with TBS-T, the fluorescence signals of the membrane were detected by an Amersham Typhoon imager. The band signal intensities corresponding to H3K36me3 were quantitated by ImageJ (*51*), and the relative signal intensities compared to that of temp42 with UTP and 3′-dATP were calculated. The mean and SD values of three independent experiments were calculated and plotted.

H3K36me3 deposition assay

Histone methylation was performed by mixing the nucleosome containing the Widom601 193 base-pair DNA (*44*), Set2, and a DY647 fluorescently-labeled RNA primer (Dharmacon) in 13.5 μL of reaction solution, and incubating the mixture at 37°C for 30 minutes. Afterward, 1.5 μL of S-adenosylmethionine (SAM) was added to the reaction solution, which was incubated for 1 hour at 37°C. The resulting reaction mixture contains 0.1 μM template nucleosome, 0.2 μM RNA, and 1.0 μM Set2, in 31 mM HEPES-KOH (pH 7.5) buffer, containing 68 mM KOAc, 0.4 μM $Zn(OAc)_2$, 0.04 mM TCEP, 2.5% glycerol, 15 mM NaCl, 0.2 mM DTT, 5 mM $MgCl_2$, 0.02 mM EDTA, 0.1 mM SAM, 0.4 mM UTP, 0.4 mM GTP, 0.4 mM CTP, and 0.4 mM 3′-dATP. A 12 μL portion of the reaction was mixed with 4 μL of 4x SDS buffer and heated at 95°C for 1 minute. The resulting samples were analyzed by western blotting, and the H3K36me3 signal was detected as described in the transcription-coupled H3K36me3 deposition assay section.

Preparation of EC-nucleosome complexes for cryo-EM analysis

The transcription reaction was conducted by mixing the template nucleosome containing H3.3 (K36M), H2B (K120Cub), and temp115 DNA, RNAPII, TFIIS, Spt4/5, Elf1, Paf1C, Spt6, Spn1, FACT, P-TEFb, CK2α, Set2, and a DY647 fluorescently-labeled RNA primer in 720 μL of reaction solution, containing S-adenosyl-L-homocysteine (SAH), UTP, GTP, CTP, and 3′-dATP, and incubated at 30°C for 75 minutes. The resulting reaction mixture contains 0.1 μM template nucleosome, 0.1 μM RNAPII, 0.1 μM TFIIS, 0.4 μM Spt4/5, 1.0 μM Elf1, 0.2 μM RNA, 0.27 μM CK2α, 0.1 μM P-TEFb, 0.4 μM Spt6, 0.5 μM Spn1, 0.4 μM Paf1C, 0.5 μM FACT, and 1.0 μM Set2, in 31 mM HEPES-KOH (pH 7.5) buffer containing 68 mM KOAc, 0.4 μM $Zn(OAc)_2$, 0.04 mM TCEP, 2.5% glycerol, 15 mM NaCl, 0.2 mM DTT, 5 mM $MgCl_2$, 0.02 mM EDTA, 0.1 mM SAH, 0.4 mM UTP, 0.4 mM GTP, 0.4 mM CTP, and 0.4 mM 3′-dATP. The resulting mixture was fractionated by the GraFix method (*52*). The sucrose gradient solution was prepared by a Gradient Master instrument (SKE), using sucrose low buffer (20 mM HEPES-KOH (pH 7.5), 50 mM KOAc, 0.2 μM $Zn(OAc)_2$, 0.1 mM TCEP, and 10% sucrose) and sucrose high buffer (20 mM HEPES-KOH (pH 7.5), 50 mM KOAc, 0.2 μM $Zn(OAc)_2$, 0.1 mM TCEP, 25% sucrose, and 0.1% glutaraldehyde). The reaction solution was applied on top of the gradient solution and centrifuged at 27,000 rpm at 4°C for 16 hours, using an SW41 rotor (Beckman Coulter). The fractions containing the EC-nucleosome complex were collected and dialyzed against 20 mM HEPES-KOH (pH 7.5) buffer, containing 20 mM KOAc, 0.2 μM $Zn(OAc)_2$, and 0.1 mM TCEP. After dialysis, the sample was concentrated by an Amicon Ultra 100K centrifugal concentrator (Millipore). For vitrification, the sample was supplemented with 0.0025% Tween-20 and applied to Quantifoil grids (copper, R1.2/1.3, 200 mesh; Quantifoil Micro Tools). The grids were glow-discharged before the sample application for 2 minutes by a PIB-10 ION Bombarder (Vacuum Device Inc.). The grids were blotted with grade 595 filter paper (Ted Pella), and then plunge-frozen in liquid ethane using a Vitrobot Mark IV (ThermoFisher) at 4°C and 100% humidity.

Cryo-EM data collection and image processing

Cryo-EM data were collected with a Krios G4 transmission electron microscope (ThermoFisher) equipped with a BioQuantum energy filter and K3 camera (Gatan). The data collection was performed using EPU (ThermoFisher), and 62325 micrographs were collected in total, with a calibrated pixel size of 0.83 Å/pix (Table S1). Subsequent image processing was accomplished with Relion 3.1 (*53*) unless otherwise specified.

The initial stages of image processing, which was aimed at creating a high-quality image stack centered at RNAPII, were performed batch-wise, similarly to that described previously (fig. S3A-C) (*20*). Briefly, the initial particle picking was performed with Relion blob picker. After preliminary 2D and 3D classifications, Topaz networks were trained with particles from several good classes, and then another round of particle picking was performed with the newly trained networks. After roughly removing bad particles with 2D and 3D classifications, all particle sets were merged, and then further purified by 2D and 3D classifications. Then, particles were re-extracted with lower binning, and the Bayesian polishing and CTF refinements were performed to improve the data quality.

To analyze the molecular structures upstream of the EC, the particles were first subjected to 3D classifications using a mask around the EC, and then particles with strong density for Spt6 were selected (fig. S3D). Then, particles were subjected to multiple local 3D classifications mainly focused on the upstream region of the EC (fig. S4), which led to the structural solution of the EC-nucleosome-Set2 complexes (figs. S6 and 7). In addition to the overall reconstruction, the upstream reconstruction and the nucleosome reconstruction were prepared, using Blush regularization implemented in Relion5 (https://www.nature.com/articles/s41592-024-02304-8).

To obtain a higher resolution map of the Spt6-Set2 interface, particles with Spt6 and Set2 were pooled and subjected to multiple 3D classifications around Spt6, using Blush regularization (fig. S5A). This resulted in the 3.06 Å reconstruction of Spt6 bound with the Set2 YKIPK peptide, and in the 3.77 Å reconstruction of Spt6 bound with Set2(AID). Also, to better visualize Set2(CD) bound to the nucleosome, upstream particles containing a suitable nucleosome and Set2(CD) were collected, and then subjected to multiple 3D classifications around Set2(CD), using Blush regularization (fig. S5B). This resulted in the 4.11 Å reconstruction of nucleosome-Set2(CD).

Model building

For the model building of the EC-nucleosome-Set2 complexes, the EC region and the nucleosome region were modeled independently, using their respective local reconstructions with higher quality. Then, the EC region and the nucleosome region were assembled into the upstream reconstructions. The model assembly was performed manually using WinCoot (*54*), ChimeraX, and ISOLDE (*55*), and then the final model was refined using Phenix (*56*).

To build the EC region including Set2(AID), the previous EC structure (PDB: 7XN7) was used as the starting model (*20*). As the map qualities of some regions (especially the RNAPII core) in the EC reconstruction were better than the previous ones, some of these regions were edited manually, and then refined with Phenix, which led to the EC model with better statistics. For both complexes (Set2$^{A}$ and Set2$^{B}$), the cryo-EM density for the DNA-RNA hybrid in the RNAPII active site was consistent with that of EC115 (fig. S8A). Spt6 and its adjacent regions were then manually edited and refined with Phenix, using the Spt6 reconstruction. The Set2 peptide region bound to Spt6 was modeled manually and refined with Phenix (fig. S8B). For the Set2-AID helical region, the model was first built with Alphafold2 (*57*), which was fit into the Spt6-Set2(AID) reconstruction, and then further refined with Phenix.

To build the nucleosome bound with Set2(CD), the nucleosomal region of the EC115 structure (PDB:7XSZ) from the previous work was used as the starting model (*20*). The initial model for Set2(CD) was created with Alphafold2, and was fit into the nucleosome-Set2(CD) reconstruction. The model was edited manually with WinCoot, ChimeraX, and ISOLDE, and then refined using Phenix. After this, the Set2(CD)-nucleosome models were slightly adjusted for Set2$^{A}$ and Set2$^{B}$, using their respective upstream and nucleosomal reconstructions.

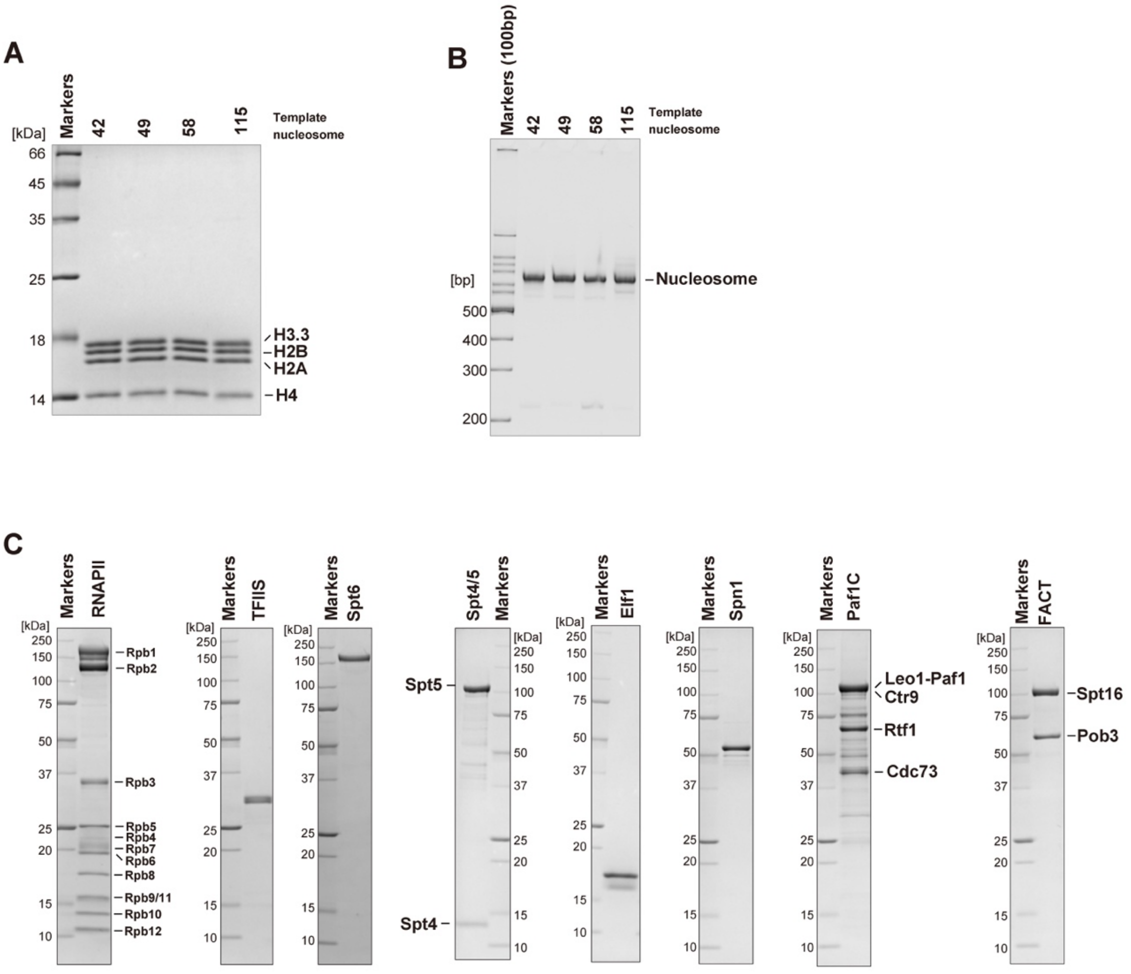

**Fig. S1. Protein preparation.** (**A**) SDS-PAGE gel of the template nucleosomes. (**B**) Native-PAGE gel of the template nucleosomes. (**C**) SDS-PAGE gels of RNAPII, transcription elongation factors, and FACT. The SDS-PAGE gels and native-PAGE gel were stained with Coomassie Brilliant Blue and ethidium bromide, respectively.

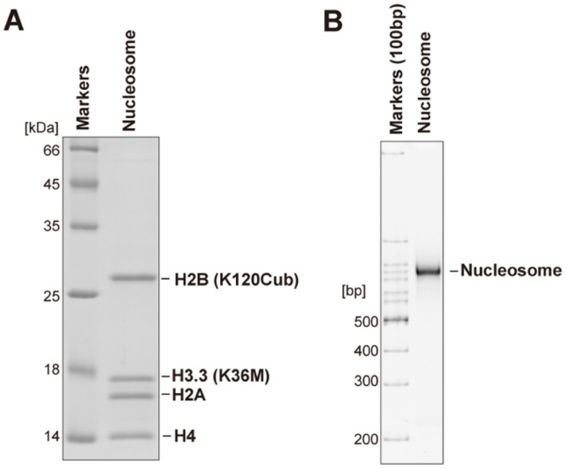

**Fig. S2. The nucleosome for the cryo-EM analysis of EC-nucleosome-Set2.** (**A**) SDS-PAGE gel of the nucleosome containing H2A, H2B (K120Cub), H3.3 (K36M), and H4. (**B**) Native-PAGE of the nucleosome. The SDS-PAGE gels and native-PAGE gel were stained with Coomassie Brilliant Blue and ethidium bromide, respectively.

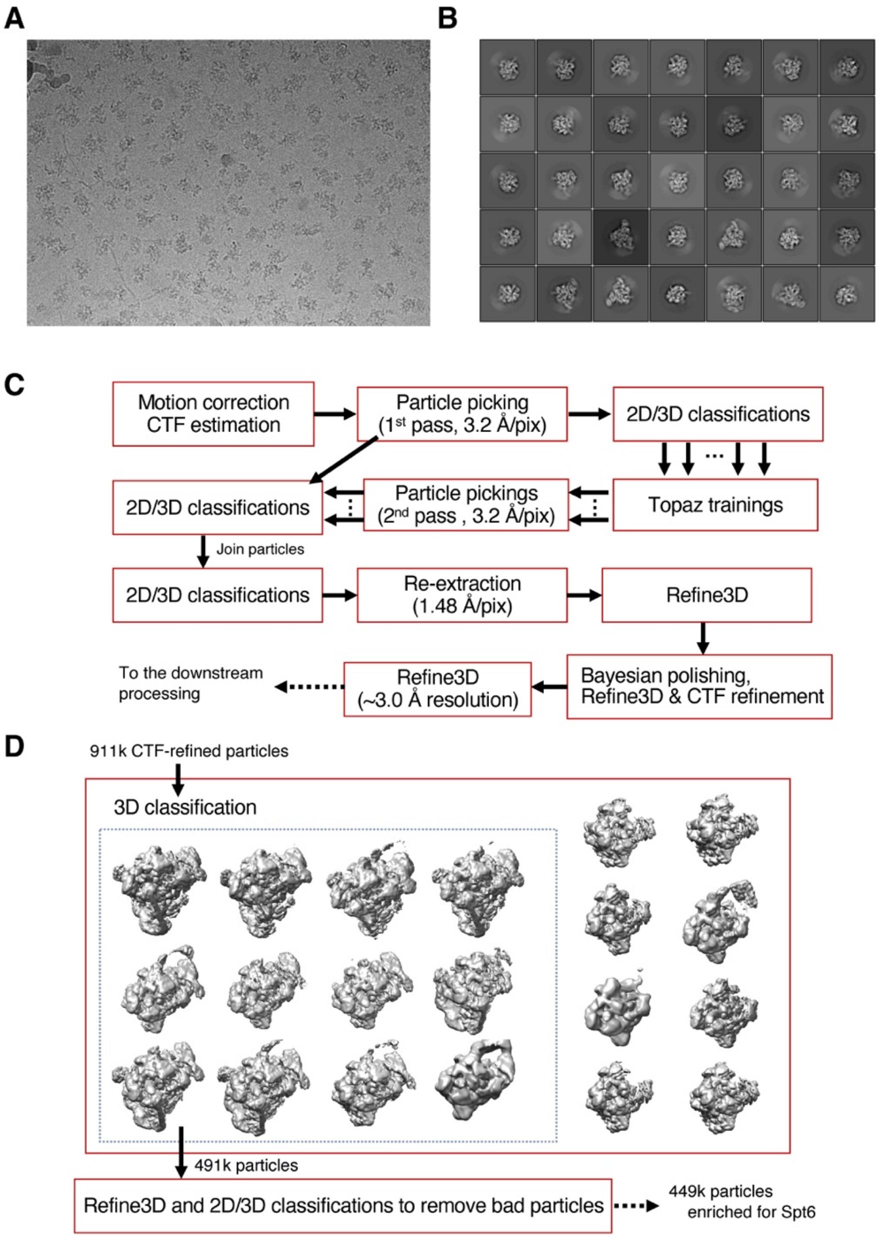

**Fig. S3. Cryo-EM data collection and initial image processing.** (**A**) Representative cryo-EM micrograph. (**B**) Representative 2D class averages from reference-free 2D classification after the Spt6-containing particles were enriched. (**C**) Representative flowchart describing the initial stage of image processing. (**D**) Representative 3D classification performed to enrich Spt6-containing particles (from one of the processing batches).

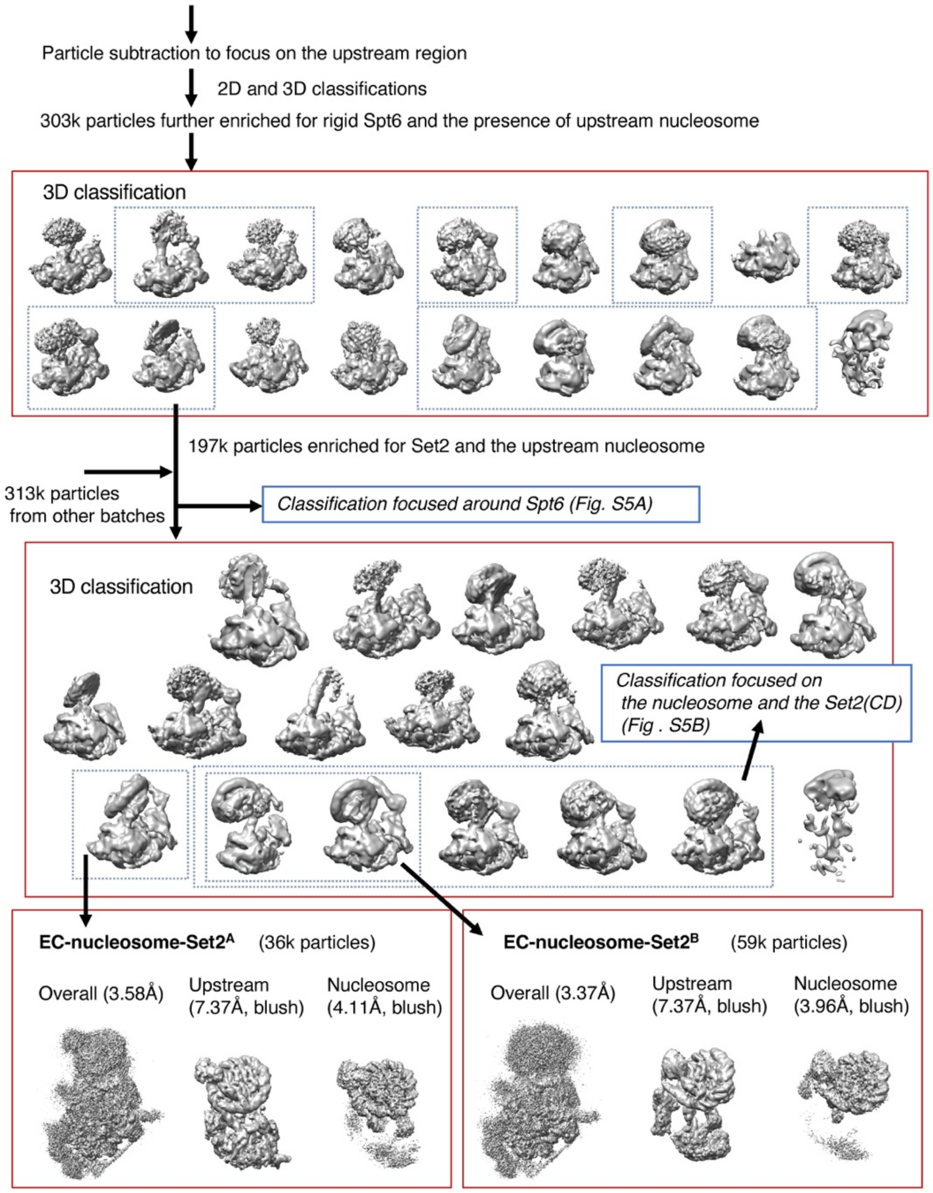

**Fig. S4. Cryo-EM data analysis leading to the EC-nucleosome-Set2 complexes.**

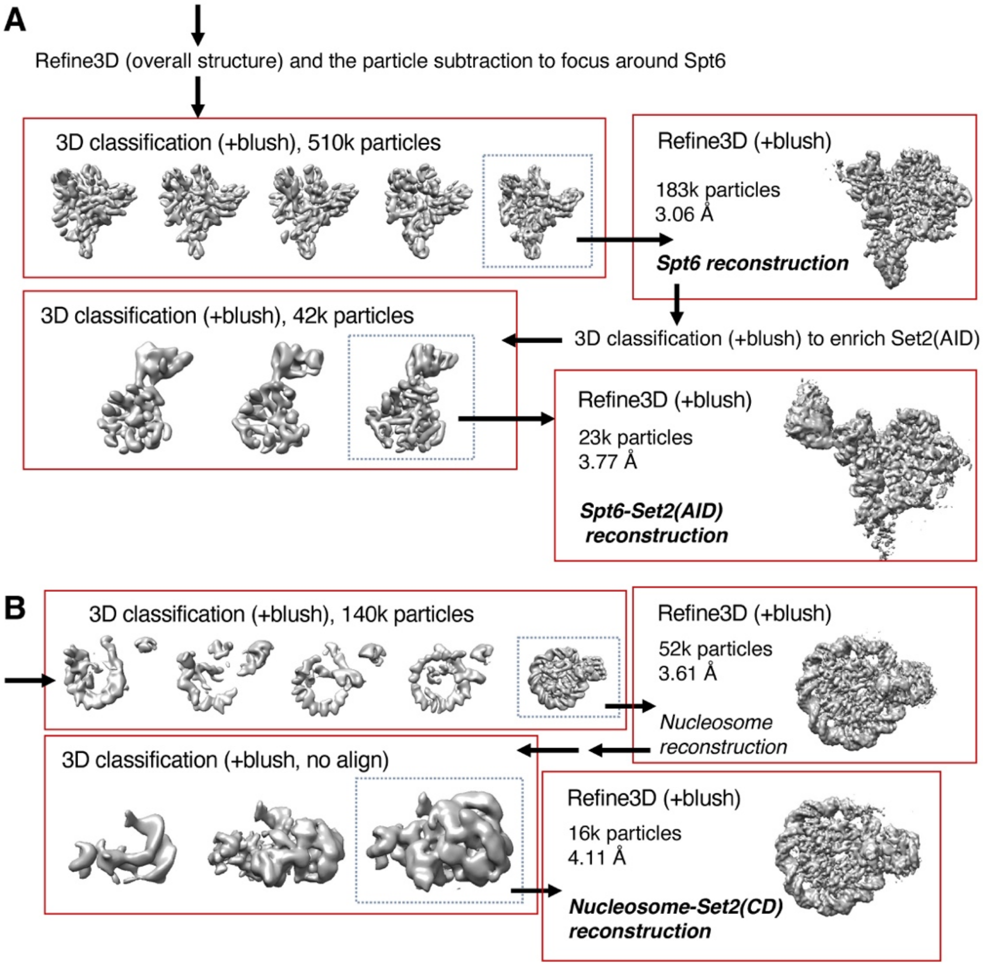

**Fig. S5. Cryo-EM data analysis leading to the local reconstructions around Set2 and Spt6.**

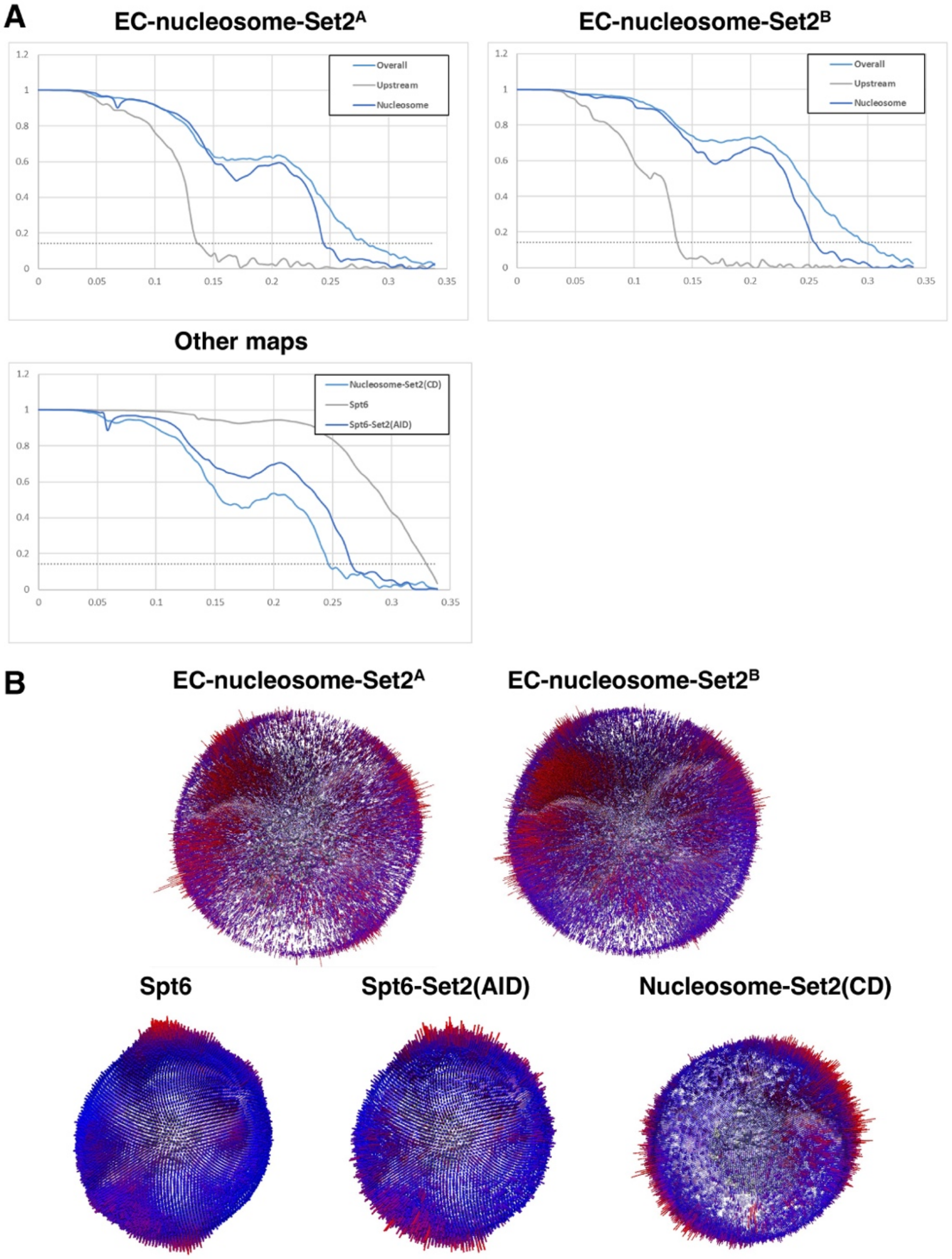

**Fig. S6. Cryo-EM statistics.** (**A**) Gold-standard Fourier shell correlation (FSC) curves of the EC-nucleosome-Set2 complexes, and (**B**) FSC curves of the local reconstructions around Set2 and Spt6. FSCs were calculated by Relion Refine3D, and dashed lines represent the FSC threshold of 0.143. (**C**) Orientation distributions of the cryo-EM reconstructions.

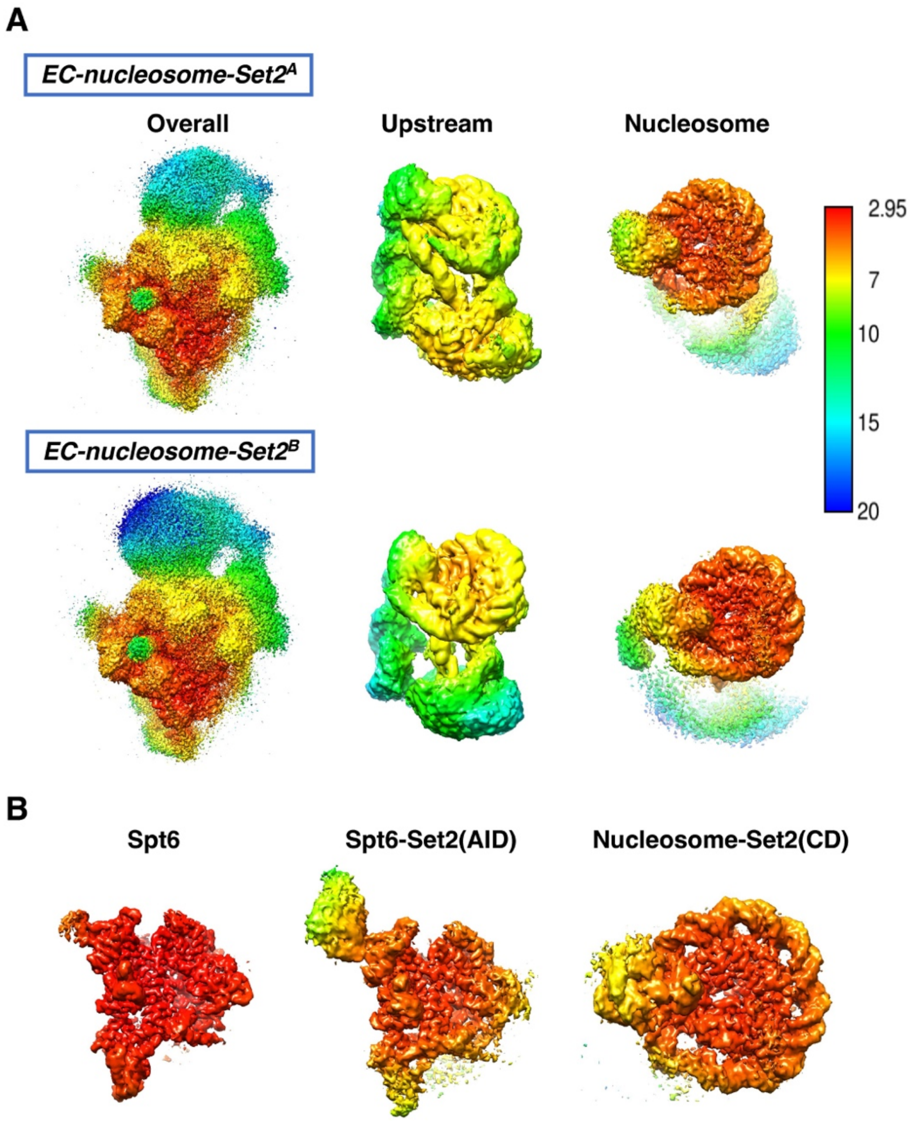

**Fig. S7. Local resolution maps of the cryo-EM reconstructions.** (**A**) Local resolutions for EC-nucleosome-Set2 complexes. (**B**) Local resolutions for the reconstructions around Set2 and Spt6. Local resolutions were calculated by Relion, with 30 Å sampling.

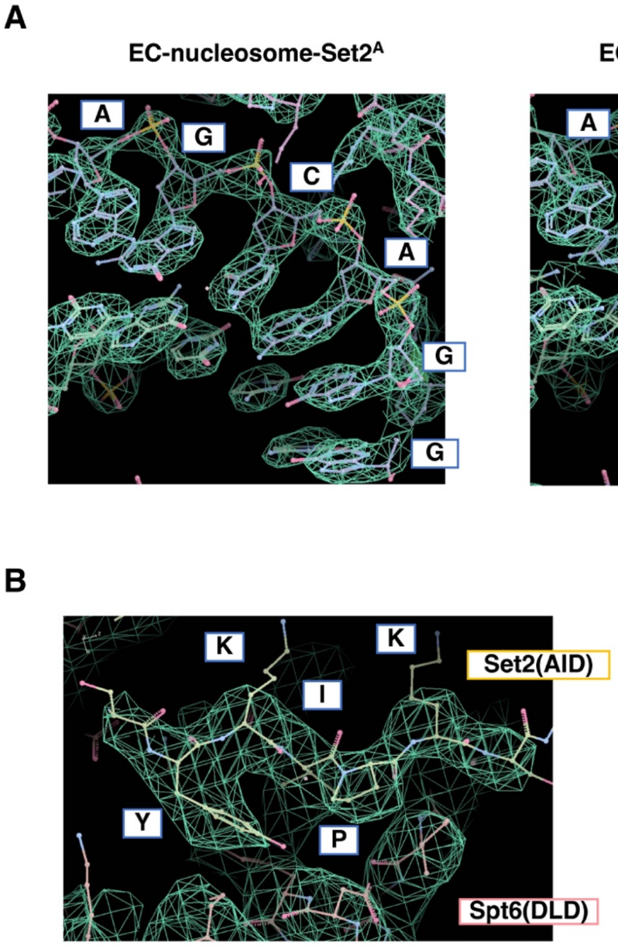

**Fig. S8. Cryo-EM densities.** (**A**) Cryo-EM maps around the EC active site from the EC-nucleosome-Set2 complexes (overall reconstruction). (**B**) Cryo-EM map around the Set2 YKIPK motif from the Spt6 reconstruction.

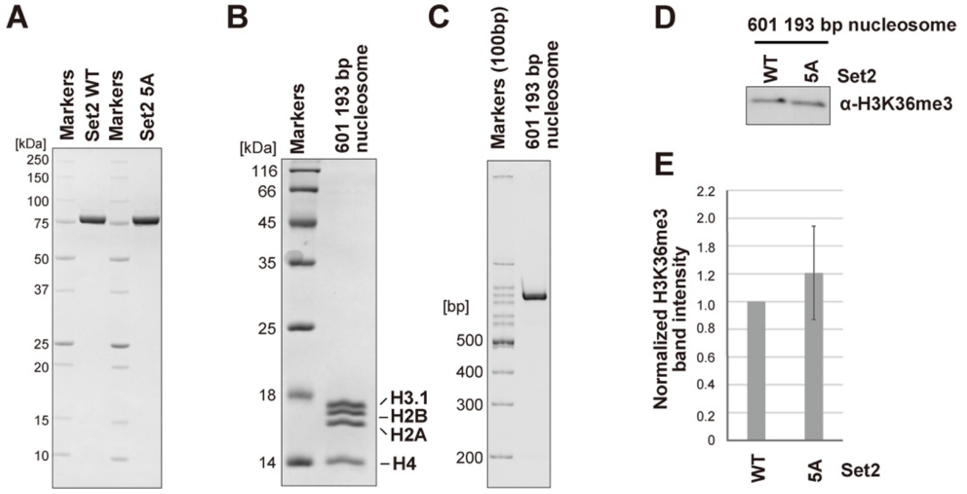

**Fig. S9. Mutational analysis of Set2.** (**A**) SDS-PAGE gel of Set2 proteins. (**B**) SDS-PAGE gels of the nucleosome containing the Widom601 193 base-pair DNA. (**C**) Native-PAGE of the nucleosome. (**D**) Western blot image of the H3K36me3 deposition assay without the EC. (**E**) Quantification of the H3K36me3 band intensities in the western blot images of four independent experiments. Relative values of the intensities compared to Set2 WT are calculated. Mean and SD are shown.

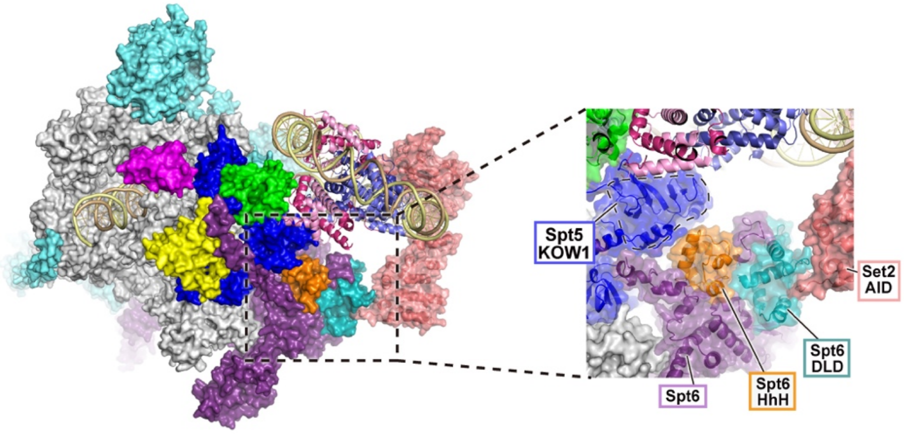

**Fig. S10. The Spt6 HhH domain.** Overall structure of EC-nucleosome-Set2$^{A}$ and close-up view of the Spt6 HhH domain. The area enclosed by a dotted rectangle is enlarged. The HhH domain is located near the Spt6 DLD domain, which is a Set2 binding site.

| Position | CDS Mutation | AA Mutation | Legacy Mutation ID | Count | Type |
|---|---|---|---|---|---|
| 2024 | c.6070C>T | p.R2024* | COSM1423518 | 2 | Substitution - Nonsense |
| 2024 | c.6071G>A | p.R2024Q | COSM3823998 | 4 | Substitution - Missense |
| 2024 | c.6071G>C | p.R2024P | COSM1045438 | 1 | Substitution - Missense |
| 2026 | c.6076C>T | p.P2026S | COSM4502193 | 1 | Substitution - Missense |
| 2026 | c.6077C>T | p.P2026L | COSM6981614 | 1 | Substitution - Missense |

**Fig. S11. SETD2 mutations in cancer.** Mutations of SETD2 2023-2027 residues in the COSMIC database (Aug. 5, 2024).

Table S1 Data collection statistics

| Sample | batch1 | batch2 | batch3 |
| --- | --- | --- | --- |
| Microscope | Krios G4 (RIKEN BDR) | Krios G4 (RIKEN BDR) | Krios G4 (RIKEN BDR) |
| Voltage (kV) | 300 | 300 | 300 |
| Detector | K3/BioQuantum | K3/BioQuantum(CDS) | K3/BioQuantum(CDS) |
| Slit width (eV) | 15 | 15 | 15 |
| Magnification | 105,000 | 105,000 | 105,000 |
| Pixel size for data collection (Å) | 0.83 | 0.83 | 0.83 |
| Total electron exposure ($e^-/Å^2$) | 61.9 | 58.9 | 56.2 |
| Exposure time (s) | 2.2 | 4.3 | 4.3 |
| Exposure rate ($e^-$/pixel/sec) | 19.4 | 9.49 | 8.98 |
| Number of frames | 48 | 48 | 48 |
| Defocus range (mm) | -1.2 to -2.0 | -1.2 to -2.0 | -1.2 to -2.0 |
| Number of collected micrographs | 13117 | 23113 | 26095 |

**Table S1. Data collection statistics.**

Table S2 Refinement and model building statistics

| | Complex[A] | Complex[B] | Spt6 | Spt6-Set2(AID) | Nucleosome-Set2(CD) |
|---|---|---|---|---|---|
| EMDB ID | EMD-XXXX | EMD-XXXX | EMD-XXXX | EMD-XXXX | EMD-XXXX |
| Number of particles | 35823 | 59019 | 183297 | 23223 | 16189 |
| Pixel size for refinement (Å) | 1.47 | 1.47 | 1.47 | 1.47 | 1.47 |
| Symmetry imposed | C1 | C1 | C1 | C1 | C1 |
| Global resolutions (Å) | | | | | |
| Overall | 3.59 | 3.37 | 3.06 | 3.77 | 4.11 |
| Upstream | 7.37 | 7.37 | - | - | - |
| Nucleosome | 4.11 | 3.96 | - | - | - |
| PDB ID | XXXX | XXXX | - | - | - |
| MolProbity score | 1.12 | 1.13 | - | - | - |
| Clash score | 3.06 | 3.08 | - | - | - |
| RMSDs | | | - | - | - |
| Bond length (Å) | 0.004 | 0.004 | - | - | - |
| Bond angle (°) | 0.706 | 0.696 | - | - | - |
| Ramachandran plot (%) | | | - | - | - |
| Outliers | 0.01 | 0.01 | - | - | - |
| Allowed | 2.08 | 2.11 | - | - | - |
| Favored | 97.91 | 97.87 | - | - | - |
| Rotamer outliers (%) | 0.06 | 0.06 | - | - | - |

**Table S2. Refinement and model building statistics.**
